# Shining Light on Osteoarthritis: Spatially Offset Raman Spectroscopy as a Window into Cartilage Health

**DOI:** 10.1101/2023.08.14.553328

**Authors:** Piyush Raj, Lintong Wu, Craig Almeida, Lauren Conway, Swati Tanwar, Jill Middendorf, Ishan Barman

## Abstract

Articular cartilage is a complex tissue, and early detection of osteoarthritis (OA) is crucial for effective treatment. However, current imaging modalities lack molecular specificity and primarily detect late-stage changes. In this study, we propose the use of Spatially Offset Raman Spectroscopy (SORS) for non-invasive, depth-dependent, and molecular-specific diagnostics of articular cartilage. We demonstrate the potential of SORS to penetrate deep layers of cartilage, providing a comprehensive understanding of disease progression. Our SORS measurements were characterized and validated through mechanical and histological techniques, revealing strong correlations between spectroscopic measurements and both Young’s modulus and depth of cartilage damage. By longitudinally monitoring enzymatically degraded condyles, we further developed a depth-dependent damage-tracking method. Our analysis revealed distinct components related to sample depth and glycosaminoglycan (GAG) changes, offering a comprehensive picture of cartilage health. Collectively, these findings highlight the potential of SORS as a valuable tool for enhancing OA management and improving patient outcomes.

## Introduction

Osteoarthritis (OA) poses a significant healthcare burden globally, affecting millions of individuals [1] [2] [3]. The disease is a slowly progressing chronic condition often characterized by degeneration of articular cartilage and eventually leads to joint pain, stiffness, and lack of mobility. Due to the slow progression of the disease, early-stage diagnosis and treatment of OA have the potential to delay or prevent the onset of severe and debilitating late-stage OA [4]. As a disease with a complex origin, understanding the molecular changes occurring in articular cartilage is crucial for early diagnosis and effective intervention. While traditional imaging modalities such as X-ray and MRI have proven valuable in visualizing large structural changes [5] [6] [7], these imaging modalities lack the ability to provide detailed molecular information, making the detection of OA at early stages challenging.

Using imaging techniques to detect molecular specificity in early-stage OA is paramount for several reasons. First, molecular alterations that precede visible morphological changes can be detected. Specifically, changes in extracellular matrix components like collagen and glycosaminoglycans (GAG) occur early in the pathogenesis of OA, even before cartilage degradation becomes evident on conventional imaging scans [8] [9] [10]. Therefore, an imaging technique with molecular specificity holds the potential to identify these early molecular markers. Secondly, molecularly specific imaging can aid in assessing treatment efficacy and understanding disease progression. By monitoring molecular changes over time, clinicians can evaluate the response to interventions at a molecular level, providing valuable insights into the effectiveness of therapeutic strategies. Moreover, molecular imaging can help identify subtypes of OA based on distinct molecular profiles, enabling tailored treatment approaches and personalized medicine. Additionally, an imaging modality with molecular specificity would offer a non-invasive means of assessing cartilage health, reducing the need for invasive diagnostic procedures such as arthroscopy or biopsy. This molecular imaging approach would provide clinicians with a valuable tool for routine monitoring of patients, facilitating timely intervention and proactive disease management.

Raman spectroscopy is a non-invasive, label-free spectroscopic technique that can address the need for molecular specificity in cartilage assessment and OA diagnostics. Raman spectroscopy is based on the phenomenon of inelastic scattering, where a tissue sample interacts with monochromatic light, resulting in characteristic energy variations between incident and scattered radiation. The resulting Raman scattering provides a unique molecular fingerprint specific to the vibrational modes, bonds, and molecules present in the tissue biochemical composition. Recently, the application of Raman spectroscopy in the investigation of rheumatologic diseases, including OA, has gained significant attention. It has been used to analyze various joint tissues such as articular cartilage, synovium, bone, and to a lesser extent, meniscus, tendons, and ligaments [11] [12] [13] [14] [15]. Researchers have identified a collection of potential optical biomarkers for OA within these tissues [16], further highlighting the potential of Raman spectroscopy in the detection of early molecular imbalances and the characterization of tissue composition in this context. However, current methods for Raman spectroscopy in cartilage assessment are limited in two critical aspects: lower penetration depth and the lack of direct correlation between spectroscopic values and damage gradation. Previous attempts to enhance signal detection from deeper cartilage layers have been restricted to a maximum depth of approximately 600 micrometers [17]. Meanwhile, human knee cartilage thickness can range from 1 to 4 mm [18]. Although most early OA changes are believed to occur near the cartilage surface, deep zone cartilage changes may also indicate early-stage OA progression [19].

In this study, we propose and demonstrate the first utilization of spatially offset Raman spectroscopy (SORS) for detecting articular cartilage degeneration. By offsetting the illumination and collection points spatially, SORS selectively probes biochemical information from different depths within the sample [20] [21] [22]. Thus, SORS provides a comprehensive assessment of depth-dependent molecular composition and surpasses the limitations of conventional Raman spectroscopy. The potential for SORS application for interrogating cartilage health, however, has been surprisingly underappreciated. Thus to establish the diagnostic potential of SORS, our study demonstrates the ability of SORS to estimate depth-dependent biochemical changes, predict biomechanical properties (notably the Young’s modulus), and estimate the degree of cartilage damage. In contrast to existing methods, our approach circumvents the complexities of multivariate curve resolution (MCR) techniques by introducing a ratiometric approach to assign spectroscopy-based glycosaminoglycan (GAG) values. This simplified method not only correlates well with mechanical measurements but also reduces hardware complexity, enabling potential translation of this method using handheld devices and faster simpler interpretation of the spectra. Moreover, longitudinal monitoring of enzymatically degraded condyle samples allows us to develop a depth-dependent damage gradation based on spectroscopic measurement. The depth-dependent damage gradation developed through longitudinal monitoring further enhances our ability to track and understand patient-specific disease progression and recovery. By employing a low-dimensional plot, we further reveal principal components dominated by sample depth and sensitivity to GAG changes. Overall, our findings emphasize the potential of SORS as an invaluable tool for advancing research and clinical diagnostics in osteoarthritis and cartilage-related disorders.

## Results and Discussion

### Spatially offset Raman spectroscopy detects signals from deeper tissue layers

Spatially offset Raman spectroscopy (SORS) has emerged as a powerful non-destructive technique for determining the subsurface chemical composition of opaque samples. Standard Raman spectroscopy techniques use inelastic light scattering of photos penetrating near the tissue surface. In contrast, SORS is based on the statistical tendency of deeper penetrating photons to undergo lateral migration from the sample’s illuminated surface in a random walk-like manner [20] [21] [22], whereas photons scattered back to the surface from shallower depths have had limited opportunity for lateral travel. Originally introduced by Matousek and co-workers [23], SORS has witnessed substantial development in recent years, with efforts focused on enhancing signal detection from deeper layers through advancements in hardware [24] [25] [26] and algorithm development [27] [28]. Biomedical applications of SORS have also been explored, such as breast cancer diagnosis [29] and assessment of bone disorders [30] [31] [32], utilizing multiple offsets. Despite the considerable research and applications of SORS, our understanding of the crucial relationship between spatial offset and depth measurement remains fragmented, primarily due to the complex interplay of absorption, scattering, anisotropy coefficients, and resultant changes in photon distribution. While some studies have investigated the probed zone extent for specific sample situations [33] [34] [35], there is a lack of generic guidance that can be universally applied across different tissue types. Recently, there have been endeavors to characterize the photon distribution using detailed Monte Carlo simulations. While these studies have offered additional insights, they have been limited to examining either a single SORS spatial offset (0.4 mm) tailored for shallow-depth skin investigations [36] or to artificially constructed phantoms composed of non-biological specimens [20]. Hence, in this study, we first sought to bridge this gap by investigating the relationship between spatial offset and depth measurement for ensuing articular cartilage assessment using SORS.

To investigate the sensitivity of measurement quantification to both the sample and the instrument, we conducted a systematic study using a custom-built Raman spectrometer system (see Materials and Methods section for details). A grocery store meat sample with a thickness of approximately 3 mm was used as the biological specimen. The experimental procedure involved acquiring on-axis Raman spectra of a plastic petri dish, followed by on-axis Raman spectra of the grocery store meat sample placed on a quartz petri dish (Figure 2a). The use of the quartz petri dish prevented spectral contamination from the plastic petri dish below, as photons originating from depths beyond 3 mm can contribute to the on-axis spectra. Distinct spectral peaks were observed in the collected Raman spectra (Figure 2b), with the plastic petri dish exhibiting peaks around 1000 cm^-1^ and the meat sample showing peaks at approximately 1440 cm^-1^. The normalized ratio of these two peaks was utilized to quantify the percentage of signal originating from the sample. This method, previously employed in [20] to characterize the signal strength from deeper layers, has hitherto not been accompanied by a comprehensive photon distribution plot.

**Figure 1:**
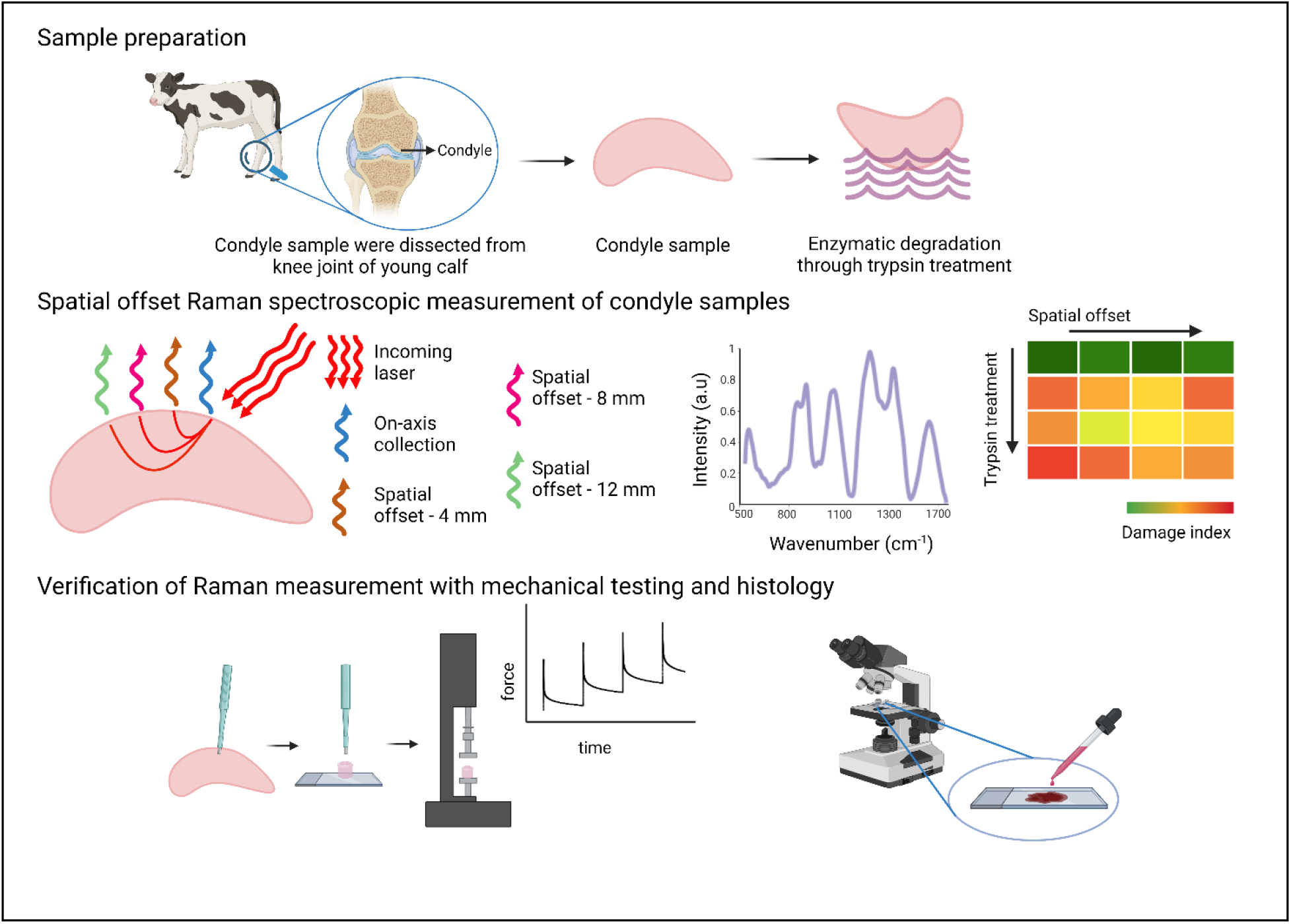
Overall schematic of study design. Articular cartilage from bovine femoral condyle was sourced. SORS measurements were performed on the untreated and treated condyle sample, and the damage gradation index was created with spectroscopic measurements. The spectroscopic measurements were cross verified with mechanical measurements and histology images.

**Figure 2:**
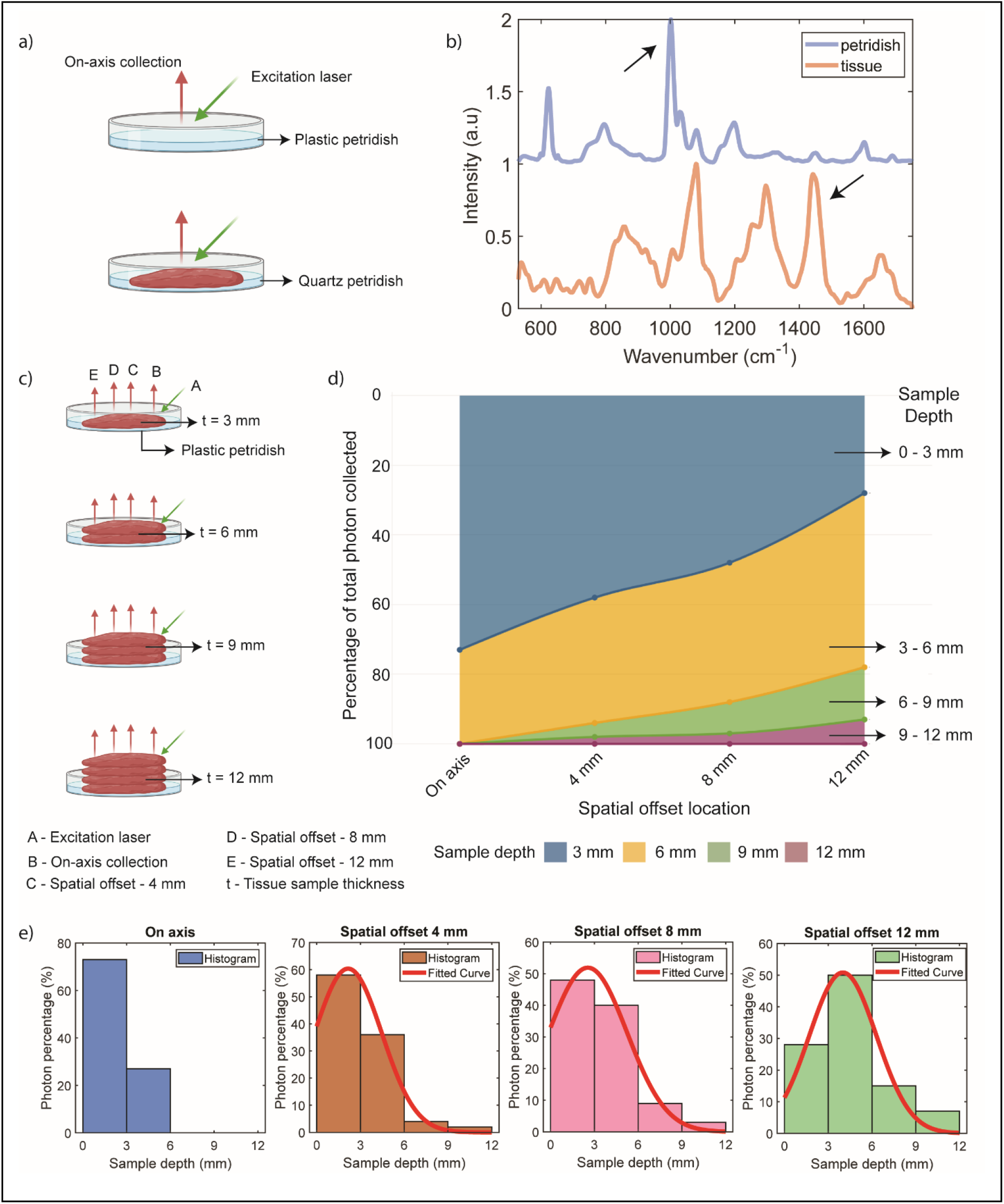
(a) Measurement schematic for collection pure spectra of the tissue sample and plastic petridish.(b) Collected Raman spectra of tissue sample and plastic petridsh highlighting the peak at 1440 cm-1 and 1000 cm-1, respectively. These peaks were used for quantifying the penetration depth of the spatial offset probe. (c) SORS measurement schematic where all the spatial offset measurements were performed for each tissue sample thickness. (d) Stacked area graph for percentage of total photon collected for different sample depth at different spatial offsets. For the case of on-axis, 73% of the photons arrive from the sample depth of 0 – 3 mm, and the remaining 27% arrive from the sample depth of 3 – 6 mm. For the case of spatial offset 4 mm, 58% of photons are coming from 0 – 3 mm depth, 34% from 3 – 6 mm depth, 4% from 6 – 9 mm depth and 4% from 9 – 12 mm depth. For spatial offset of 8 mm, 48% of photons are coming from 0 – 3 mm depth, 38% from 3 – 6 mm depth, 9% from 6 – 9 mm depth and 5% from 9 – 12 mm depth. Finally for spatial offset of 12 mm, we have 28% of photons coming from 0 – 3 mm depth, 50% from 3 – 6 mm depth, 15% from 6 – 9 mm depth and 7% from 9 – 12 mm depth. (e) Gaussian fitted curve (except for on-axis) to quantify the correlation between spatial offset and sample depth.

To explore the relationship between spatial offset and sample depth, multiple spatial offset measurements were taken for each sample thickness (Figure 2c). The signal strength originating from the plastic petri dish below was quantified as a percentage. A stacked area graph (Figure 2d) illustrates the photon distribution from the biological specimen. For the on-axis collection, 73% of photons originated from depths within 0-3 mm, with 27% originating from depths within 3-6 mm. No photons were collected below 6 mm depth in the on-axis configuration. For a spatial offset of 4 mm, the majority of photons (58%) originated from depths within 0-3 mm, with 34% from 3-6 mm, 4% from 6-9 mm, and 4% from 9-12 mm. Similarly, with a spatial offset of 8 mm, the predominant depth range for photon contribution was 2-4 mm, with 48% originating from 0-3 mm, 38% from 3-6 mm, 9% from 6-9 mm, and 5% from 9-12 mm. With a spatial offset of 12 mm, the main depth range shifted to 3-6 mm, with 28% of photons originating from 0-3 mm, 50% from 3-6 mm, 15% from 6-9 mm, and 7% from 9-12 mm. Further details on the quantification methodology can be found in Section I of the supplementary information. The correlation between spatial offset and sample depth was quantified by fitting a Gaussian curve to represent the percentage of photons originating from each quantized depth (Figure 2e), barring the on-axis case where the presence of only two data points precluded such analysis.

These experimental measurements reveal a distinct correlation between spatial offset and the depth to which photons penetrate in biological specimens. The findings consistently demonstrate that increasing the spatial offset allows for the detection of signals originating from deeper layers which is consistent with previous measurements [20]. In addition to that, we calibrated our measurement system against biological specimens which had not been systematically explored before. Crucially, the distribution of photons concerning spatial offset and depth was quantitatively characterized, enhancing the interpretability of SORS measurements.

### Spatial offset Raman spectroscopy accurately predicts depth-dependent cartilage degradation

Articular cartilage possesses a unique structure and composition that endows it with exceptional mechanical performance. It comprises a network of type-II collagen (COL) fibrils, providing structural support and tensile and shear strength [37] [38]. Additionally, the cartilage matrix contains negatively charged sulfated glycosaminoglycans (GAGs), which contribute to its compressive properties and facilitate water retention within the tissue [39]. Loss of GAG is a common occurrence during early-stage osteoarthritis. This loss of GAG leads to weakened cartilage, which could cause other degenerative changes including cellular damage and inflammation. Consequently, GAG content serves as an important marker for cartilage health assessment. Owing to its importance, Raman spectroscopic measurements have been extensively employed to investigate GAG degradation and cartilage organization [13] [16] [40] [41] [17]. However, most studies have qualitatively assessed GAG depletion by comparing samples with pristine cartilage or stained histopathology images. To the best of our knowledge, only one study has correlated Raman measurements at different treatment points with indentation elastic modulus, albeit with limited depth penetration capabilities of 600 um [17]. It is also worth noting that indentation elastic modulus focuses primarily on the surface response and may not provide a complete picture of the cartilage’s mechanical behavior. Given that GAG depletion during osteoarthritis (OA) initially occurs in the upper portion of the middle zone and progresses to the deep zones [18], a depth-based assessment of GAG is crucial to comprehensively evaluate cartilage health.

With the aim of exploring the capability of SORS to detect differences in signals from deeper layers, we conducted a series of experiments on femoral condylar cartilage (Figure 3). The cartilage was divided into four different groups, each comprising three condyles subjected to trypsin treatment for varying durations: 0 hours, 3 hours, 6 hours, and 9 hours. The treatment was conducted using a specially designed fixture that ensured only the top surface of the samples came into contact with trypsin [Refer to section III in supplementary information]. To assess the depth-dependent changes, SORS measurements were performed using specific spatial offsets of 0 mm, 4 mm, 8 mm, and 12 mm. These spatial offsets were chosen based on the previous section, which provided estimates of the photon distribution from different depth sections. In all the spatial offset measurements, a reduction in the signal corresponding to the Raman peak associated with GAG was observed as the duration of directional trypsin treatment increased (highlighted in figure 3(c) and 3(d)). Notably, the maximum signal reduction was observed in the on-axis measurements, which corresponded to the top layer experiencing the most enzymatic degradation. Conversely, at the maximum spatial offset of 12 mm, the signal reduction caused by increased spatial offset was minimal as the deeper layers experienced less degradation. The other two spatial offset measurements (4 mm and 8 mm) exhibited varying degrees of GAG signal reduction, reflecting the degree of directional enzymatic degradation. A detailed band assignment of the Raman peaks in our cartilage measurements can be found in the supplementary information.

**Figure 3:**
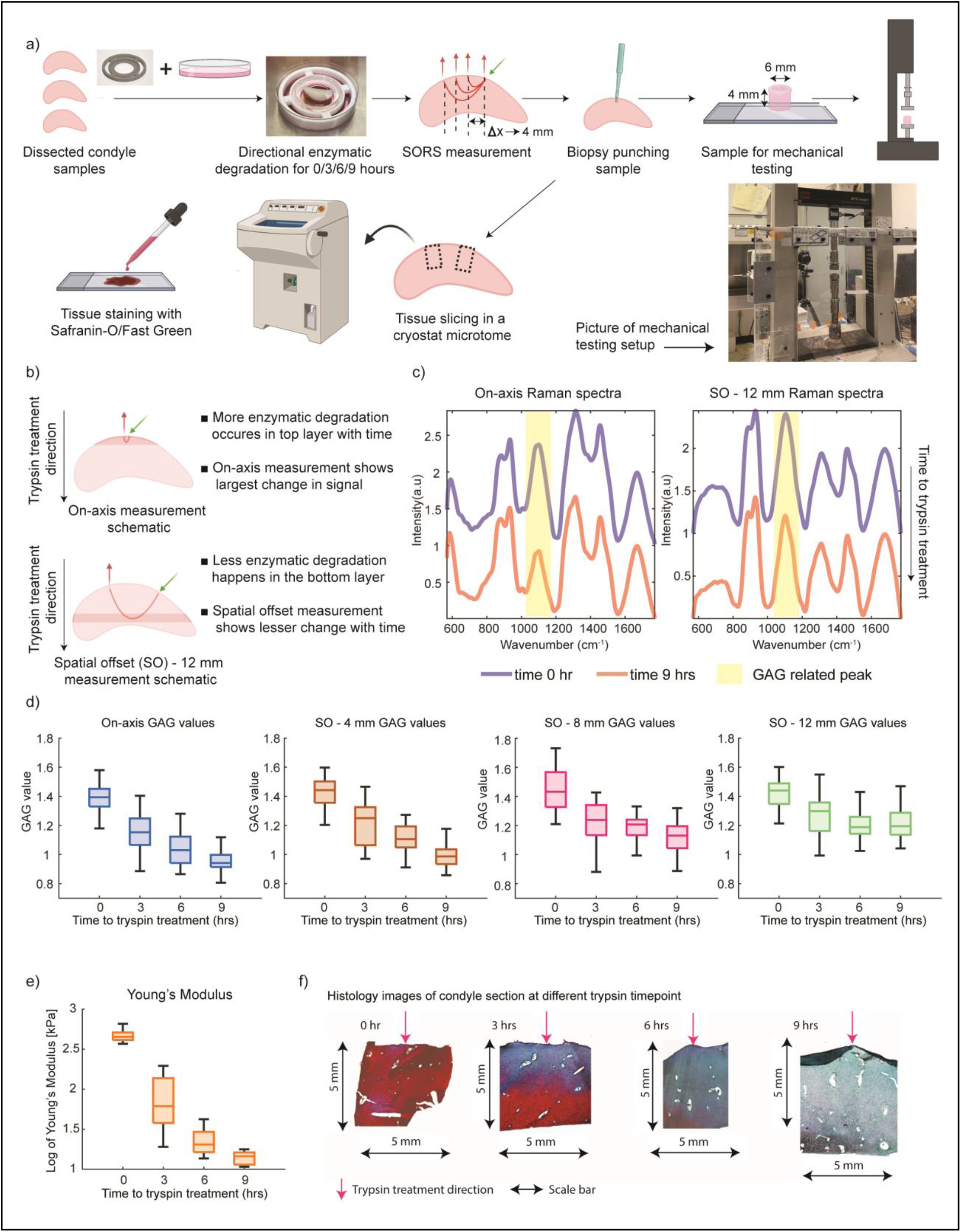
(a) Study design for establishing correlation between non-invasive SORS measurement with mechanical and histology measurements. (b) Figure explaining the reason for differential GAG signal at different spatial offset at different timepoint. (c) Mean Raman spectra after background subtraction and normalization across three condyle samples for on-axis and spatial offset 12 mm collection for pristine condyle sample and 9-hour trypsin-treated sample. (d) GAG value for different spatial offset at different trypsin treatment timepoint highlighting that more degradation is occurring in the top layer and less degradation happens in the bottom layer. (e) Log of Young’s modulus value of condyle plugs about 6 mm in diameter and 4 mm in depth at different trypsin treatment timepoint. (f) Safranin-O/Fast Green histology-stained images of condyle section at different trypsin treatment timepoint verifying the findings of the spectroscopic measurement.

After completing the Raman spectral acquisition, the samples underwent mechanical measurements and histology. The entire workflow is presented in Figure 3(a). In short, two to three 6 mm diameter biopsy punches were taken from each condyle and cut to a height of approximately 4 mm. Then, samples underwent unconfined compression testing. Samples were subjected to a series of five 5% strain steps until 25% compressive strain was reached. Between each strain step the samples were allowed to stress relax and reach a stress equilibrium. The Young’s modulus was directly measured from these experiments and showed a decreasing trend with increasing trypsin treatment (Figure 3(e)). In contrast to indentation measurements, which are sensitive to boundary conditions, unconfined compression testing provides accurate measures of isolated cartilage samples. This technique measures the average mechanical changes throughout the entire depth of the cartilage tissue, capturing the contributions of changes that may occur in both the superficial and middle zones.

After mechanical testing, samples were fixed in formalin, paraffin embedded, and sectioned onto charged glass slides. Then Safranin-O/Fast Green histology was performed to qualitatively assess GAG content in each sample. Increasing the duration of trypsin treatment increased the loss of GAG content (Figure 3(f)). In 3-hour trypsin samples, GAG content was typically degraded up to 2.5 mm. In both 6-hour and 9-hour trypsin samples GAG content was depleted to the full depth of the sectioned sample 5 mm. These histology differences can explain the observed differences in the Raman probe measurements. Specifically, the lack of GAG signal differences between 6- and 9-hour trypsin groups may be associated with the fact that most of this signal comes from between 3 and 6 mm.

In our study, we established a direct correspondence between the reduction in GAG signal measured by Raman spectroscopy and the Young’s modulus measured through mechanical testing. Figure 4 illustrates the strong correlation between the on-axis Raman measurement and the logarithm of Young’s modulus, with a correlation coefficient of 0.93. Similarly, a high correlation of 0.9 was observed for the spatial offset of 4 mm. It is noteworthy that even though the Young’s modulus exhibited significant heterogeneity within the same trypsin timepoint, these changes were consistently captured by Raman spectroscopy and correlated with the mechanical measurements. However, for spatial offsets of 8 mm and 12 mm, where a substantial portion of the signal originated from deeper layers, the correlation was weaker. In these cases, histology images (Figure 3(e)) provided valuable insights by confirming the increase in GAG depletion to depths below 2 mm, 4 mm, and 5 mm at 3 hours, 6 hours, and 9 hours of trypsin treatment, respectively.

**Figure 4:**
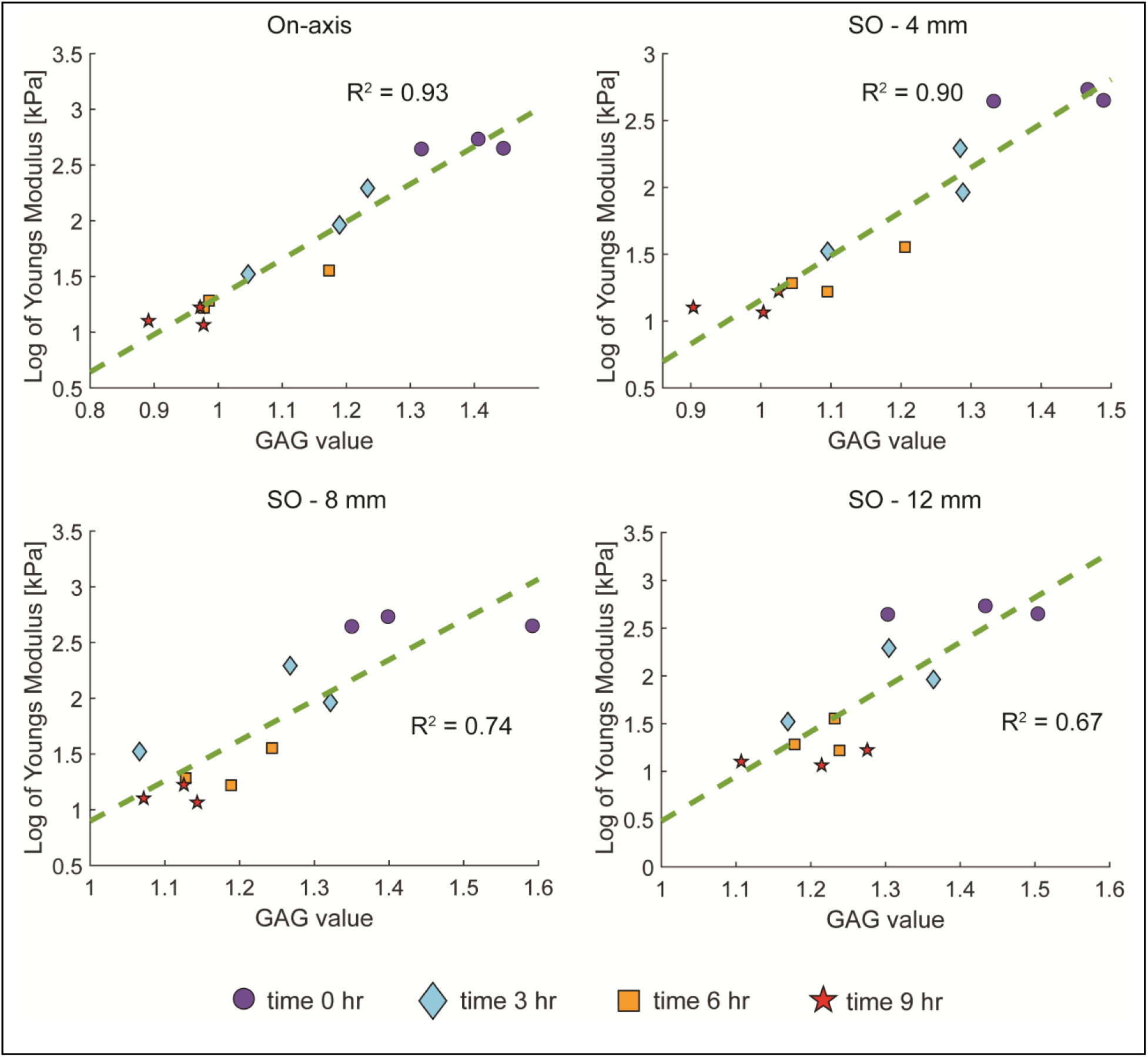
Correlation between GAG values measured with Raman spectroscopic measurements and log of Young’s modulus (kPa) value measured through compression testing.

Notably, prior studies have demonstrated that molecular alterations occur early in the pathogenesis of osteoarthritis (OA), preceding the detectable degradation of cartilage on conventional imaging scans [42] [43]. These molecular changes involve key components of the extracellular matrix, such as collagen and glycosaminoglycans (GAGs), which play crucial roles in maintaining the structural integrity and mechanical properties of cartilage. In line with this notion, our experiments use enzymatic degradation, a well-established method to simulate a diseased state in articular cartilage. This degradation method showed minimal changes in the physical shape and visual appearance of the samples between the pristine and enzymatically degraded states. This suggests that the early molecular modifications which are detected by Raman measurements [44] has the potential for preventive monitoring. Identifying and targeting these molecular changes in the early stages could lead to the development of improved biomolecular assays for early detection and monitoring of disease progression, potentially enabling interventions before irreversible structural damage occurs.

Furthermore, we would like to highlight the analysis methodology employed in this study, which resulted in the high correlation observed. Rather than relying on complex multivariate analysis techniques, we employed a simple and effective approach. The Raman spectrum was normalized with the water peak, and the normalized signal at 1080 cm^-1^ was used as the GAG value. This straightforward method yielded a significantly higher correlation compared to previously reported techniques [13] [17]. This strong correlation is of great importance as it enables the prediction of Young’s modulus values solely from Raman spectroscopic measurements. Additionally, the comprehensive series of SORS measurements provides valuable information for estimating the depth of degradation with a few millimeters accuracy. The simplicity of this approach not only opens up possibilities for developing miniaturized Raman systems but also reduces the burden on hardware by utilizing filters instead of gratings, resulting in higher photon efficiency [45]. Moreover, the assessment process becomes instantaneous, and the interpretability of the data becomes more straightforward, facilitating its translation into clinical applications.

### Longitudinal Assessment of Cartilage Health Using Spatial Offset Raman Spectroscopy

Building upon the successful application of the SORS method to predict Young’s modulus of cartilage and estimate the depth of degradation, we sought to design a study that enables the longitudinal monitoring of the same condyle sample across multiple time points. In our previous experiments, we conducted tests on multiple condyles, introducing inherent variability due to biological differences and the nature of trypsin diffusion, despite maintaining consistent trypsin treatment times and methods. To overcome these challenges and accurately track the enzymatic degradation process, we implemented a longitudinal monitoring approach. This involved assessing the same condyle sample at different time points of trypsin treatment. The complete workflow of this longitudinal monitoring study is depicted in Figure 5(a).

**Figure 5:**
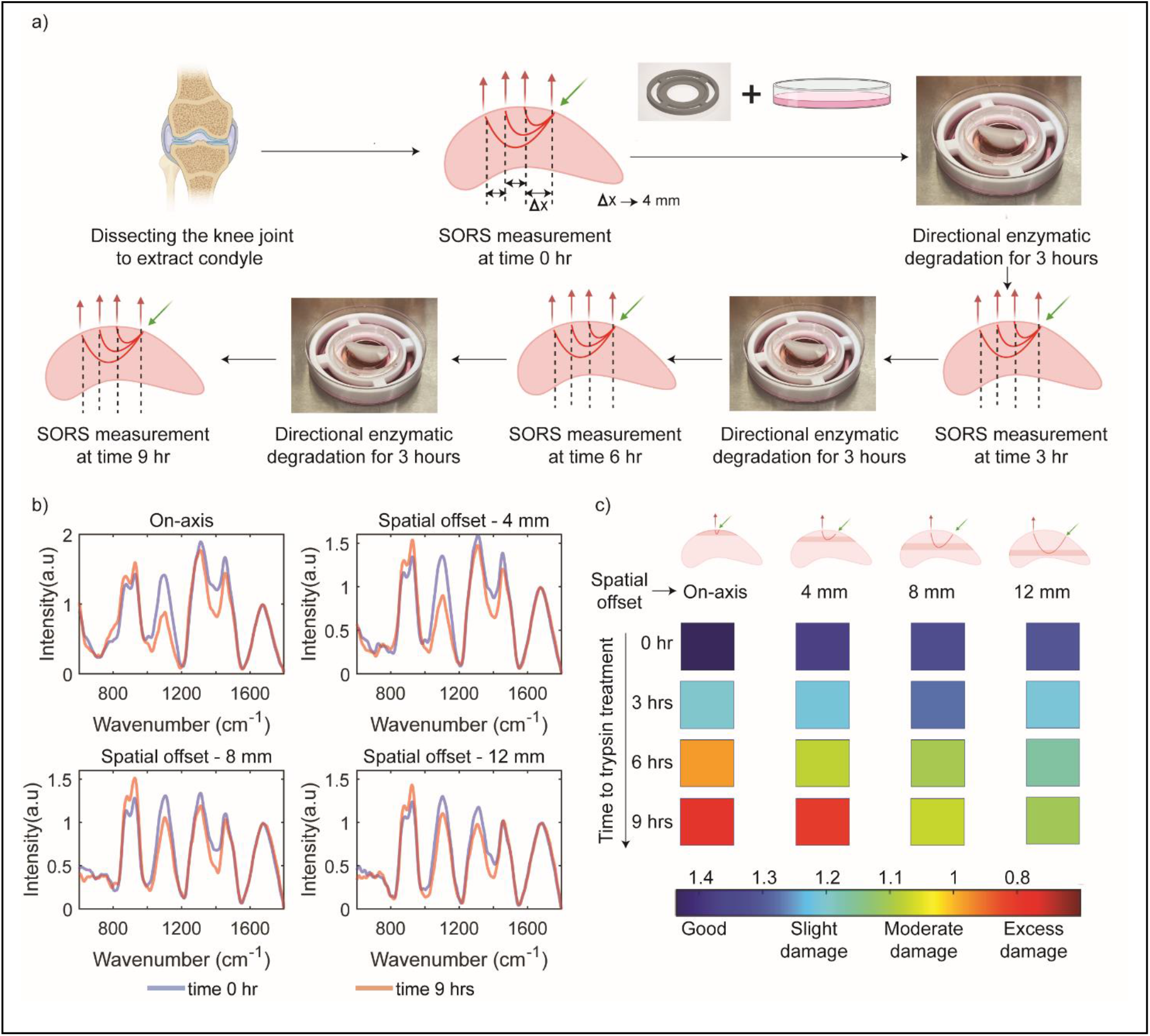
(a) Measurement schematic for longitudinal monitoring of condyle sample. (b) Mean Raman spectra for all spatial offsets before trypsin treatment and after 9-hours of trypsin-treatment. (c) Figure highlighting the possibility of depth-based damage gradation with Raman spectroscopic measurement.

Upon initial inspection of the Raman spectra, similar trends to the previous discrete case are observed. The largest difference in GAG signal strength due to increasing trypsin treatment times was observed in the on-axis spectra. This difference decreased with increasing spatial offsets of 4 mm, 8 mm, and 12 mm. Figure 5 (b) displays the Raman spectra for time 0 hours and time 9 hours at different spatial offsets. The gradual progression of GAG degradation at different time points, probed at different depths by the varying spatial offsets, is depicted in Figure 5 (c).

When longitudinal data is combined with our calibrated probe depth data, the depth dependent progression of GAG depletion can be identified. We can associate spatial offset of Raman probe with penetration depth in the sample through the calibration measurement performed in the earlier section (Figure 2). The on-axis measurement primarily probes the top layer (0-2 mm) of the sample, aligning with the reported maximum degradation in the top layer for directional trypsin treatment (Figure 3). Consequently, the GAG signal exhibits the lowest strength for the 9-hour treatment, with intermediate values progressing from 3 hours to 6 hours and finally 9 hours. A similar decline in GAG values is observed for the spatial offset of 4 mm, which probes a depth of approximately 1-3 mm. At 6 hours, a slightly different color indicative of cartilage health is observed compared to the on-axis measurement, as the damage has not extended deeper than 4 mm in the condyle sample. For spatial offsets of 8 mm and 12 mm, which probe depths of approximately 2-4 mm and 4 mm onwards, respectively, a decrease in GAG signal corresponding to GAG depletion is observed. At the 3-hour trypsin treatment time point, negligible GAG degradation is observed below 3 mm; however, a reduction in GAG signal is still observed in the spectroscopic measurement. This phenomenon occurs because not all photons originate from the assigned depth when assigning sample depth based on spatial offset measurements. The photon distribution follows a left-skewed curve with respect to sample depth [46] [see section II in supplementary information where we have provided details of Monte Carlo simulation of photon distribution]. As a result, even at larger spatial offsets, there is a small percentage of photons originating from the top layer, leading to some GAG signal reduction at larger spatial offsets during the 3-hour trypsin treatment.

The use of a single condyle allowed for the tracking of gradual spectral changes on a low-dimensional principal component (PC) plot. To conduct this analysis, all spectra from all offsets and degradation treatment times were combined into a single data matrix, and principal component analysis (PCA) was performed. PCA is a statistical technique used to reduce the dimensionality of data and identify the most important patterns or variations in the dataset. The scores of the first and second PCs were then plotted. Upon careful examination of the PC plots, it is evident that the PC1 component is primarily influenced by spectral variations originating from measurements at different depths, while the PC2 component reflects the degradation of GAG. Together, these plots enable the tracking of sample-specific, depth-based cartilage degradation at different depths and time points. Figure 6(a) illustrates that measurements performed at different spatial offsets cluster together along the PC1 axis. The multivariate nature of the spectral measurements captures changes in the spectra when measurements are conducted at various offsets, probing different depths of the sample. In Figure 6(b), the PC2 axis serves as a proxy for GAG signal strength, revealing the progression of damage at different sample depths. The highest damage is observed for the on-axis measurement, with a gradual decline at larger spatial offsets. Notably, for a spatial offset of 12 mm, the measurement points nearly merge, highlighting the minimal damage occurring at a depth of 5 mm and beyond. These PC plots provide valuable insights into the depth-dependent changes and can aid in the objective and early assessment of cartilage degradation.

**Figure 6:**
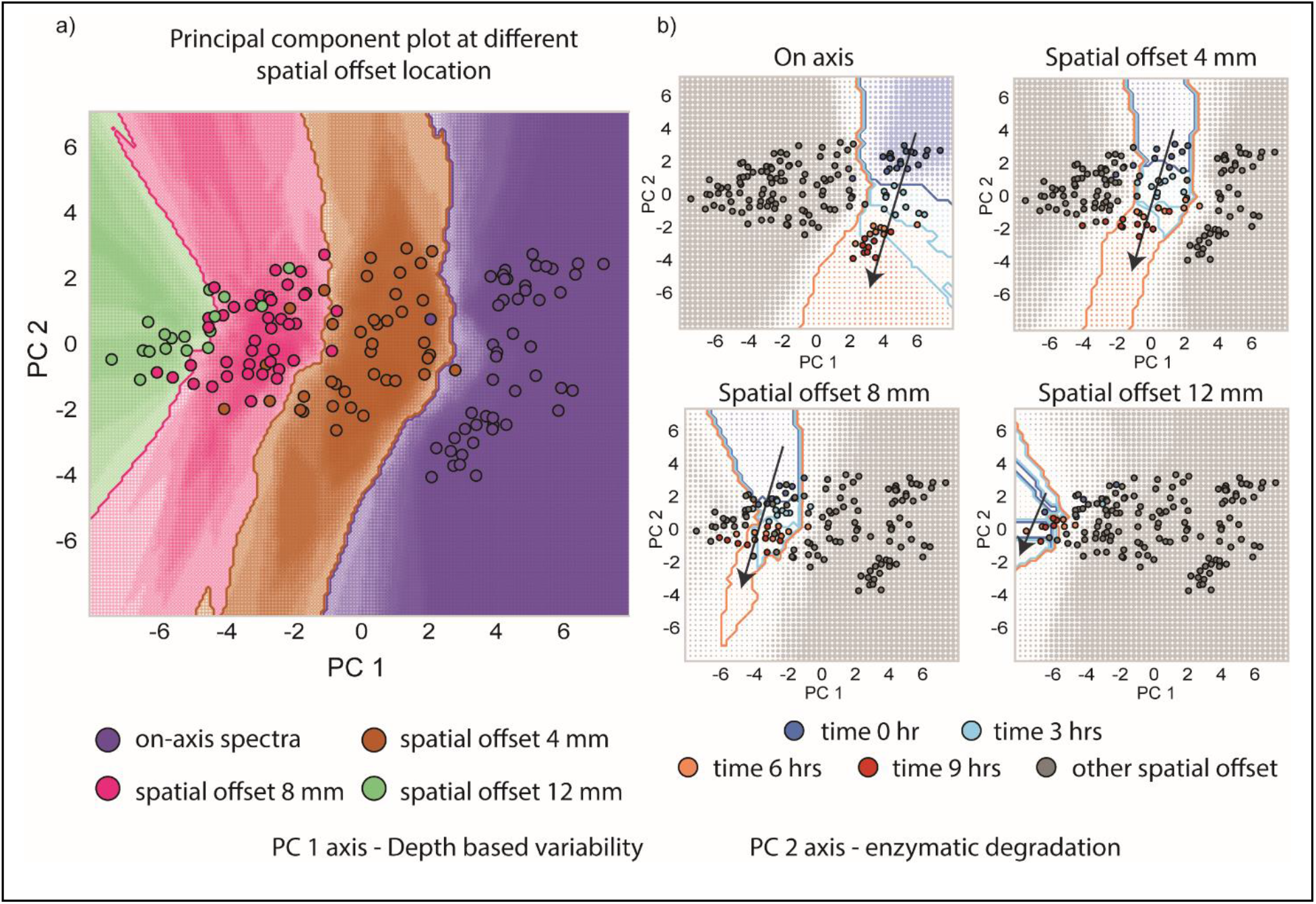
(a) Principal component plot of all the spectra collected during longitudinal monitoring. PC 1 and PC 2 scores are plotted which correspond to depth-based variability and enzymatic degradation respectively. Different spatial offset spectra are clustered at different values along PC 1 axis. (b) Each PC plot highlights a set of spatial offset points that are colored according to the different trypsin treatment times. The progression of patient specific damage at different depths can be tracked with these PC plots.

## Conclusion

In conclusion, spatially offset Raman spectroscopy (SORS) emerges as a powerful tool for non-invasive and depth-dependent assessment of articular cartilage. This study demonstrates the potential of SORS in capturing molecular changes that occur early in the pathogenesis of osteoarthritis (OA), even before structural and mechanical changes to the tissue. By accurately predicting cartilage health and providing molecular-specific information, SORS offers new opportunities for early detection and monitoring of OA. The correlation between spectroscopic measurements and mechanical properties highlights the value of SORS in evaluating cartilage health and understanding the underlying molecular alterations. Furthermore, the longitudinal monitoring approach employed in this study opens the door for monitoring degeneration induced in animal models of OA and monitoring patient-specific damage progression. By combining SORS measurements with principal component analysis, the study successfully tracks molecular changes at different depths and time points, offering a comprehensive assessment of cartilage degradation. Collectively, these findings signify a significant step towards the advancement of molecular-specific spectroscopic measurements for early diagnosis and personalized treatment of articular cartilage diseases. With further refinement and expansion of SORS techniques, including human samples and in vivo measurements, this imaging modality holds great promise for improving osteoarthritis management and patient outcomes. Future research should explore the potential of SORS in other musculoskeletal disorders and continue to push the boundaries of molecular-specific imaging in the field. Ultimately, SORS has the potential to revolutionize our approach to cartilage assessment, paving the way for more targeted and effective interventions in the field of musculoskeletal health.

## Materials and Methods

Cartilage from the femoral condyles of neonatal bovine knees was collected (Animal Technologies, Tyler, TX).

### Trypsin treatment process

The dissected condyle sample was placed on the 3D printed fixture designed according to the dimension of the condyle, and the 3D printed holder was placed in 100 mm × 20 mm style petridish. 13 ml 0.25% trypsin-EDTA (1×) solution was added into the dish, and the tip region of the curved surface of the condyle sample was submerged in 0.25% trypsin-EDTA (1×) solution. The trypsin treatment was carried out in the CO2 incubator at 37 °C with 5% CO_2_ concentration, and the trypsin treatment durations are 3, 6, and 9 hours. After trypsin treatment, the condyle sample was washed with 1× PBS solution three times.

### Raman spectroscopic measurement

The Raman measurements were conducted using a custom-built Raman probe system designed for excitation and collection of Raman signals. The excitation probe utilized a 300-micron core fiber for laser delivery and incorporated a low-pass filter to ensure excitation at 830 nm wavelength. The collection probe consisted of concentrically arranged fibers along the circumference of a 2 mm diameter probe. It was equipped with a 2 mm diameter truncated ball lens and a 1 mm thick magnesium fluoride window at the probe tip. Both probes were manufactured by EmVision LLC (https://emvisionllc.com). The spatial offset was created by placing the two probes on a custom-designed fixture (see section IV in supplementary information). The angle between the probe was 30 degrees.

The excitation and collection probes were connected to a portable clinical Raman system, which included an 830-nm diode laser (Process Instruments, maximum power: 500 mW), a spectrograph (Holospec f/1.8i, Kaiser Optical Systems), and a thermoelectrically-cooled CCD camera (PIXIS 400BR, Princeton Instruments). To maintain consistent laser power, the laser was operated at an appropriate current value to deliver a fixed power of 70 mW at the sample.

Raman data acquisition parameters varied depending on the measurement configuration. For on-axis measurements, an integration time of 5 seconds with 3 accumulations was used. Spatial offset measurements at 4 mm utilized an integration time of 10 seconds with 3 accumulations. For spatial offset measurements at 8 mm, an integration time of 15 seconds with 3 accumulations was employed. Lastly, spatial offset measurements at 12 mm utilized an integration time of 25 seconds with 5 accumulations.

### Raman spectral analysis

The wavenumber calibration of the Raman system was performed using acetaminophen spectra acquired by the same system. The spectral analysis focused on the fingerprint region between 570 cm^-1^ and 1,770 cm^-1^. To enhance the quality of the acquired Raman spectra, several pre-processing steps were applied. These included fifth-order polynomial-based background subtraction, median filtering, and vector normalization. The background subtraction reduced fluorescence contributions, the median filtering mitigated cosmic rays, and the vector normalization compensated for laser power variations. In the normalized spectra, the GAG value was directly derived from the signal at 1080 cm^-1^.

For the principal component analysis (PCA), the “pca” function in MATLAB was employed to analyze the spectral data and extract the principal components. In addition, a high-density PCA plot with a decision boundary was generated using the Python programming language. The scikit-learn library was utilized to train a K-nearest neighbor (KNN) classifier on the given classes and PC scores data. A custom grid size was predefined, and the grid points were predicted using the trained KNN classifier. The color of each grid point represented the predicted class, while the size of the point indicated the probability of the classification. The plots were created using the matplotlib library. A detailed implementation of this approach can be found at the following link: https://www.tvhahn.com/posts/beautiful-plots-decision-boundary/

### Mechanical measurement details

Condyles from SORS measurements were then subjected to classic unconfined compression analysis [47] [48]. To prepare samples for mechanical testing, a 6 mm diameter biopsy punch was used to create two to three 6 mm diameter plugs from each condyle. The plugs were then trimmed to a thickness of 4 mm. The thickness was checked using a caliper. Then, plugs were loaded onto an MTS Insight 5 SL testing machine (MTS, Eden Prairie, MN) with the articular surface (superficial zone) facing up. The upper platen was brought into full contact with the cartilage. Throughout testing, samples were immersed in Phosphate buffered saline (PBS) and were progressively strained in five steps of 5% each to a total strain of 25%. The strain was ramped at 0.1 mm/s [49]. After each strain step, the samples were allowed to stress relax for at least 5 minutes or until the normal load changed by less than 0.01N/min. The Young’s Modulus of each cartilage plus was calculated directly from the equilibrium stress-strain data.

Correlations between Young’s modulus and the GAG concentration were completed using a linear least squares analysis.

### Histology methods and steps

After mechanical testing, Safranin-O/Fast Green histology staining was completed to qualitatively measure GAG depletion. First, cartilage plugs were fixed in 10% buffered formalin for 24 hours, followed by ethanol. Then, plugs were frozen in an optical cutting temperature (OCT) medium and cryo-sectioned into 12 μm slices onto oxygen plasma-cleaned quartz slides. Tissue sections were stained with modified Wiegert’s iron hematoxylin for 5 minutes and further treated with a 1% acid alcohol solution to remove excess stain. Sections were then stained with 0.02% fast green for 1 minute and rinsed with 1% acetic acid solution. These sections were stained with 1% safranin O for 30 minutes. Stained sections were then dehydrated with increasing concentration of ethanol followed by xylene treatment. Imaging was carried out immediately after staining using PrimeHisto XE histology slide scanner.

## Acknowledgment

IB would like to acknowledge the support of the National Institute of General Medical Sciences (1R35GM149272). JM would like to acknowledge support from startup funding provided by Johns Hopkins University.

