## Supplementary information for "Shining Light on Osteoarthritis: Spatially Offset Raman Spectroscopy as a Window into Cartilage Health"

#### I. Method on how photon distribution plot was made:

To understand how the photon distribution was calculated, we first define the signal strength. 100% plastic petridish signal is defined as ratio of 1000 $cm^{-1}$ to 1440 $cm^{-1}$ for pure plastic petridish spectra and 0% plastic petridish signal is defined as ratio of 1000 $cm^{-1}$ to 1440 $cm^{-1}$ for pure meat sample spectra.

We now take example of on-axis measurement with different sample thickness. For each measurement, we know that all of the signal is coming from 0 mm and beyond since the probes are pointed downward. Therefore,

| | On - axis |
|---|---|
| 0 mm and beyond | 100 % photons |

Now, we put a 3 mm thick meat sample on plastic petridish and perform on-axis measurement. We find that 27% of signal came from plastic petridish. This means that 27% of the signal came from 3 mm and beyond depth.

| | On - axis |
|---|---|
| 0 mm and beyond | 100 % photons |
| 3 mm and beyond | 27% photons |

Next, we put another layer of meat sample making it 6 mm meat layer sample. We see that 100% of signal came from meat sample and 0% came from plastic petridish. The same thing happens when a total of 9 mm and 12 mm thick tissue sample was placed on petridish for on-axis measurement. Therefore, we have the following:

| | On - axis |
|---|---|
| 0 mm and beyond | 100 % photons |
| 3 mm and beyond | 27% photons |
| 6 mm and beyond | 0% photons |

| | |
|---|---|
| 9 mm and beyond | 0% photons |
| 12 mm and beyond | 0% photons |

The same process was repeated for all spatial offset measurements which results in the following matrix:

| | On axis | Spatial offset 4 mm | Spatial offset 8 mm | Spatial offset 12 mm |
|---|---|---|---|---|
| 0 mm and beyond | 100 | 100 | 100 | 100 |
| 3 mm and beyond | 27 | 42 | 52 | 72 |
| 6 mm and beyond | 0 | 8 | 14 | 22 |
| 9 mm and beyond | 0 | 4 | 5 | 7 |
| 12 mm and beyond | 0 | 0 | 0 | 0 |

To quantify the photons coming from each layer, the photon percentage was subtracted from the nearest row:

| | On axis | Spatial offset 4 mm | Spatial offset 8 mm | Spatial offset 12 mm |
|---|---|---|---|---|
| 0 – 3 mm | 73 [100 – 27] | 58 [100 – 42] | 48 [100 – 52] | 28 [100 – 72] |
| 3 - 6 mm | 27 [27 – 0] | 34 [42 – 8] | 38 [52 – 14] | 50 [72 – 22] |
| 6 – 9 mm | - | 4 [8 – 4] | 9 [14 – 9] | 15 [22 – 7] |
| 9 – 12 mm | - | 4 [4 – 0] | 5[5 – 0] | 7 [7 – 0] |

### II. Spatial offset Raman spectroscopy simulation for different offset

Monte Carlo methods was used to simulate the following spatial offset Raman spectroscopy simulation in MATLAB. The following simulation parameters were considered:

- Simple one-layer model
  - Layer optical property ($u_a$ = 0.1 $cm^{-1}$, $u_s$ = 62.5 $cm^{-1}$ , g = 0.9)

- Simulation boundary = 1.2 cm cube
- Simulation step = 0.003 cm
- Refractive Index matched (No Fresnel reflection considered)
- Beam type
  - Gaussian, Near-field – Top hat, Far-field – Gaussian, Diameter = 0.06 cm
- Collector
  - Diameter = 0.04 cm, NA = 0.35

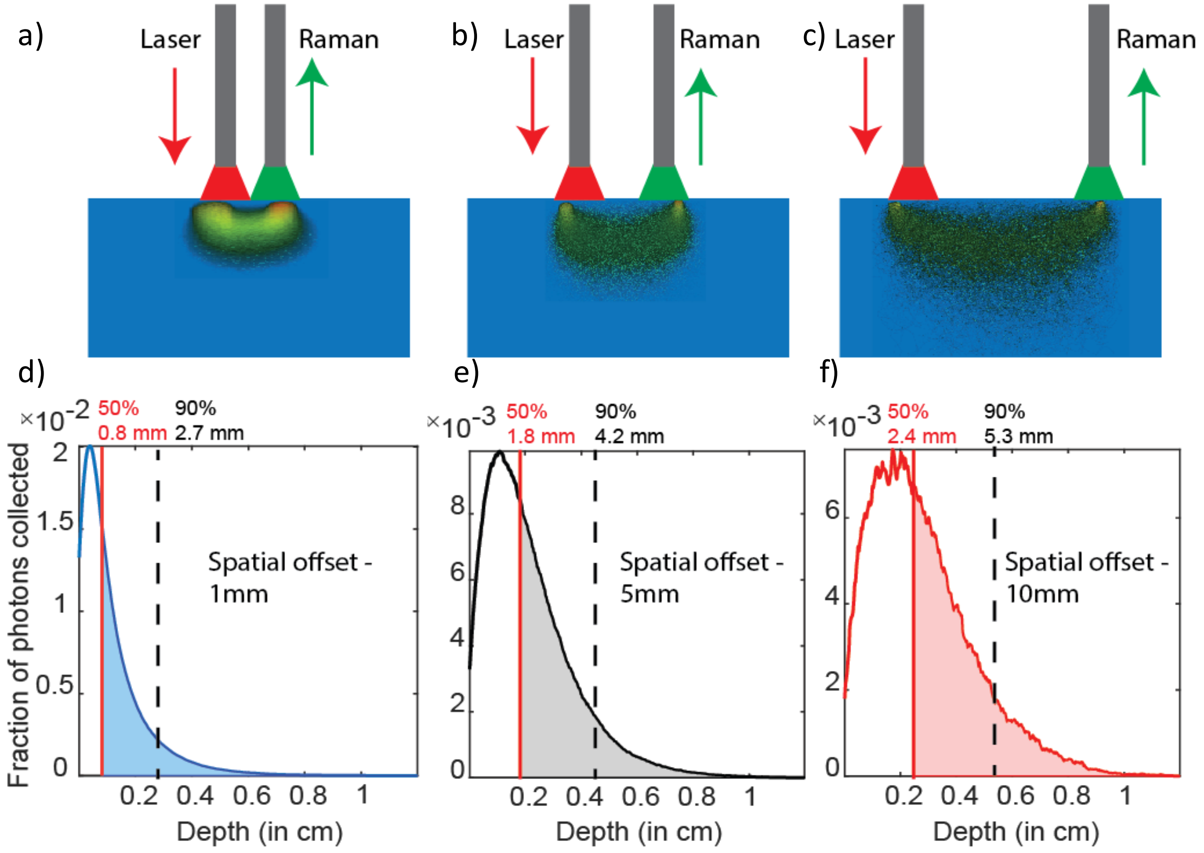


**Figure 2.1**: In figures (a) and (d), the spatial offset is 1 mm. The photon trajectory in figure (a) clearly shows that most of the photons reaching the collector are probing the shallow layer compared to other ones. In this configuration, 50% of the photons reaching the collector come from layer 0.8 mm and below, and 90% of photons come from range 0 to 2.7 mm. In figures (b) and (e), the spatial offset is 5 mm. In this configuration, 50% of the photons reaching the collector come from layer 1.8 mm and below, and 90% of photons come from range 0 to 4.2 mm. In figures (c) and (f), the spatial offset is 10 mm. In this configuration, 50% of the photons reaching the collector come from layer 2.4 mm and below, and 90% of photons are coming from range 0 to 5.3 mm.

### III. Fixture design for directional enzymatic degradation

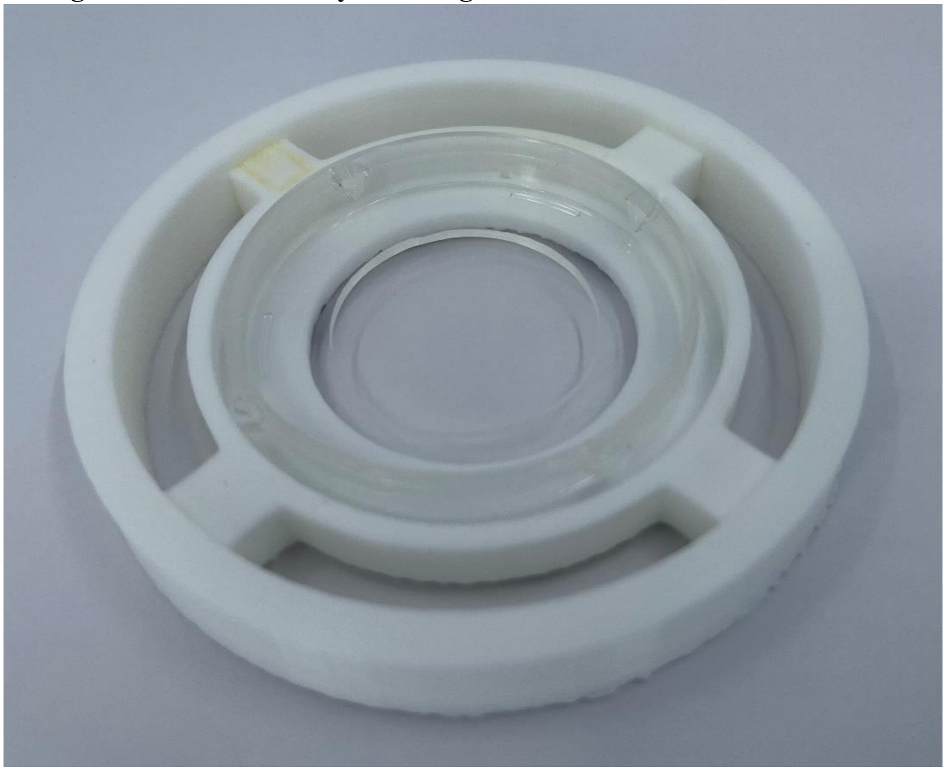

**Figure 3.1:** 3D-printed fixture design for directional trypsin treatment of condyle

### IV. Picture of SORS setup

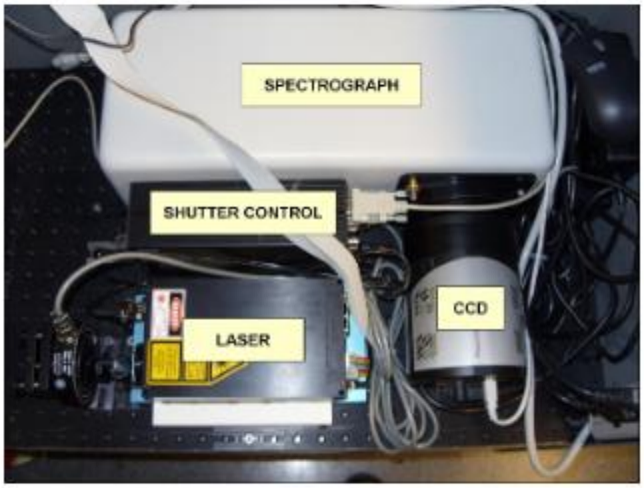


**Figure 4.1:** Major instrument component of Raman spectroscopy on the optical breadboard

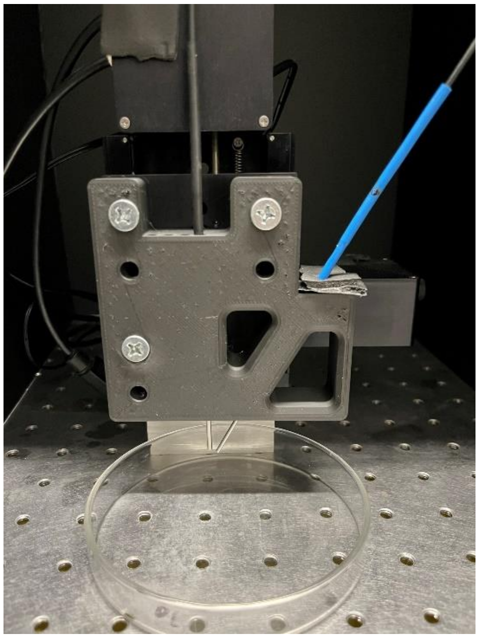

**Figure 4.2:** Sample measurement stage with SORS probes mounted. Excitaiton probe is blue in color and collection probe is black in color.

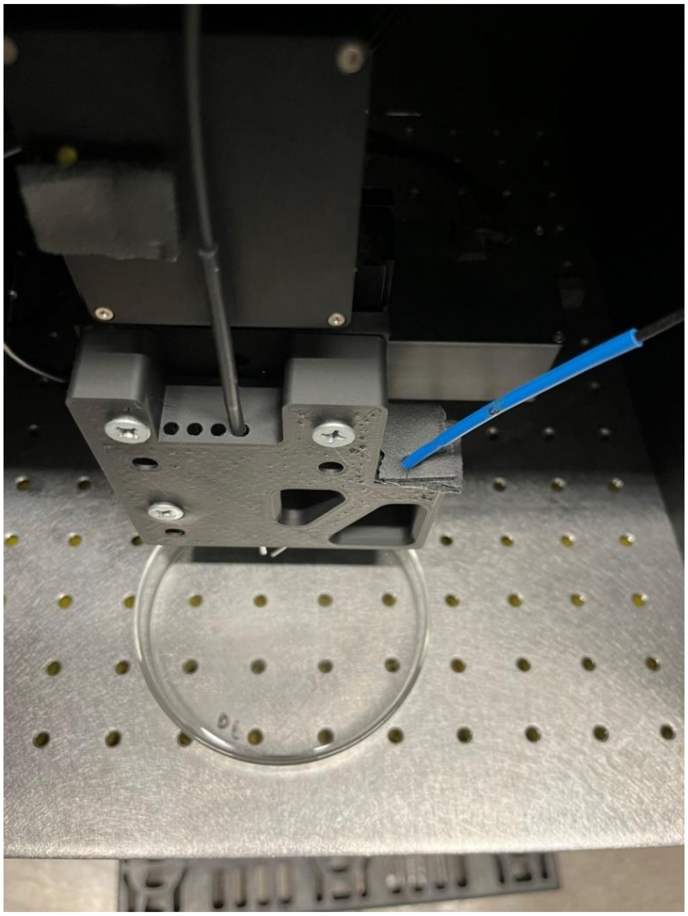

**Figure 4.3:** Top view of the SORS fixture showing multiple offset location for collection fiber probe

V. **Band assignment of cartilage Raman spectra**

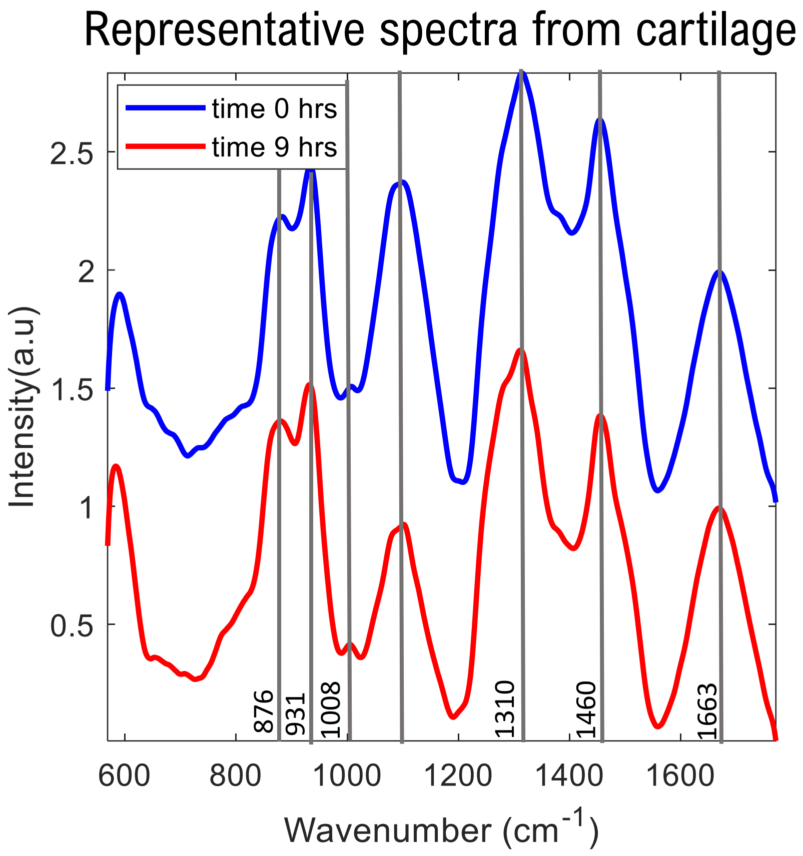


**Figure 5.1:** The spectra shown is collected at spatial offset 4 mm at two trypsin treatment timepoint – 0 hours [no treatment] and 9 hours.

| **Raman peak** | **Assigned bond/molecule** | **Component** | **Ref.** |
|---|---|---|---|
| 876 $cm^{-1}$ | C–C stretching | Collagen | [1] [2] |
| 931 $cm^{-1}$ | Symmetric stretching: C–C protein backbone | Collagen | [1] [2] [3] [4] |
| 1008 $cm^{-1}$ | Phenylalanine (C-C aromatic ring) | Proteins | [4] |
| 1080 $cm^{-1}$ | O –$SO_3^-$ symmetric stretching | Sulphated GAGs, PGs, Aggrecan | [5] [6] |
| 1310 $cm^{-1}$ | Hydrated amide C=O | Proteins | [6] [7] |

| | | | |
|---|---|---|---|
| | and N–H (α-helix) | | |
| 1460 $cm^{-1}$ | $-CH_2$ and Amide I | Lipid | [8] |
| 1663 $cm^{-1}$ | 0-H-0 | Water | [9] |
